# PAF15 stabilizes PCNA on DNA and slows down its sliding dynamics

**DOI:** 10.1101/2025.11.15.688598

**Authors:** Daehyung Kim, Amaia Gonzalez-Magaña, Athokpam Langlen Chanu, Gayun Bu, Muhammad Tehseen, Cherlhyun Jeong, Jae-Hyung Jeon, Samir M. Hamdan, Alfredo De Biasio, Francisco J. Blanco, Jong-Bong Lee

**Author notes:** Correspondence to: A.D.B., F.J.B. or J.-B.L.

## Abstract

The PCNA-associated factor 15 (PAF15) protein is essential for human genome stability by regulating DNA replication and repair through its interaction with proliferating cell nuclear antigen (PCNA). Despite its significance, the mechanistic details of the PAF15-PCNA interaction remain insufficiently understood. Here, we reveal how PAF15 influences the sliding dynamics and function of PCNA. Using single-molecule diffusion analysis on a DNA skybridge platform, we show that PAF15 binding stabilizes PCNA in its DNA-bound state, reduces its diffusion rate along the duplex, restricts accessibility to PCNA PIP-box binding sites, and regulates PCNA loading and unloading on DNA via replication factor C (RFC). Our results are consistent with its coordinating role in both high-fidelity replication and lesion bypass. These findings establish PAF15 as a key regulator of PCNA function, serving as a mobile platform for DNA-editing enzymes and thereby influencing genome maintenance.

## Introduction

The DNA-sliding clamp proliferating cell nuclear antigen (PCNA) plays a critical role in eukaryotic DNA replication and repair^1^. Human PCNA forms a homotrimeric ring-shaped structure that encircles the DNA duplex and anchors DNA polymerases, thus increasing their processivity, the number of incorporated nucleotides between enzyme association and dissociation events^2^. By interacting with various DNA-editing enzymes and regulatory proteins, PCNA serves as a versatile platform that coordinates DNA synthesis and repair processes to maintain genomic stability^3^. In human cells, PCNA is predominantly loaded on DNA by the heteropentameric ATPase Replication Factor C (RFC)^4^. While RFC also exhibits limited unloading activity, efficient PCNA unloading is primarily mediated by the ATAD5-RFC complex, where the RFC1 subunit is replaced by the ATAD5 one^5^.

PCNA’s ability to recruit different proteins is largely mediated by the PCNA-interacting protein (PIP) motif, present in various binding partners^6^, which bind to the PIP-binding sites on the front face of the ring. This and other PIP-like motifs^7^ allow proteins to dock on PCNA. As there are three identical PIP-binding sites, the so-called toolbelt model proposes that more than one enzyme can bind simultaneously to the same PCNA ring^8^, enabling the timely recruitment of specific factors needed for DNA synthesis and damage tolerance. This is especially relevant for Okazaki fragment maturation, as the lagging-strand DNA polymerase, flap-endonuclease, and ligase must act sequentially. This model is supported by cryoEM structures of PCNA/DNA/Polymerase-δ/FEN1^9^ and PCNA/DNA/Lig1/FEN1^10^ complexes. Additionally, recent cryo-EM structures of endogenously purified Okazaki fragment maturation complexes revealed FEN1 and Ribonuclease H2 (RNaseH2) simultaneously bound to PCNA, supporting the notion that toolbelt assemblies form in vivo^11^. However, PCNA’s interactions must be carefully regulated to avoid competition or conflicts among its binding partners, especially during replication stress or DNA damage^12,13^.

PCNA-associated factor 15 (PAF15, also known as PCLAF15) is one of PCNA’s key regulatory partners^14^, and its association with PCNA is emerging as an important regulatory mechanism in DNA replication and repair^15,16^. PAF15 is an intrinsically disordered protein^17^ that uniquely binds PCNA: the central region containing the PIP motif binds on the PIP-binding sites of PCNA, its chain threading through the hole, and the N-terminal and C-terminal tails emerging disordered at opposite sides of the ring, accessible for interactions and posttranslational modifications^18,19^ (**Figure 1A**). The N-terminal region binds DNA with micromolar affinity, and it becomes doubly monoubiquitinated to recruit DNA Methyltransferase 1 (DNMT1) to the replication fork^20–22^. The free PCNA ring can simultaneously bind three PAF15 chains, but the crystal structure of a PAF15 fragment bound to PCNA with a DNA duplex inside the ring suggests that more than two PAF15 chains will create steric clashes^23^. This structure showed that PAF15 limits the sliding surface available for DNA. This interaction, however, may need to be released for efficient DNA lesion bypass by switching from replicative to translesion synthesis (TLS) polymerases. Most of PCNA-associated PAF15 is ubiquitinated, and its degradation by the proteasome facilitates the polymerase switch^24^.

**Figure 1.**
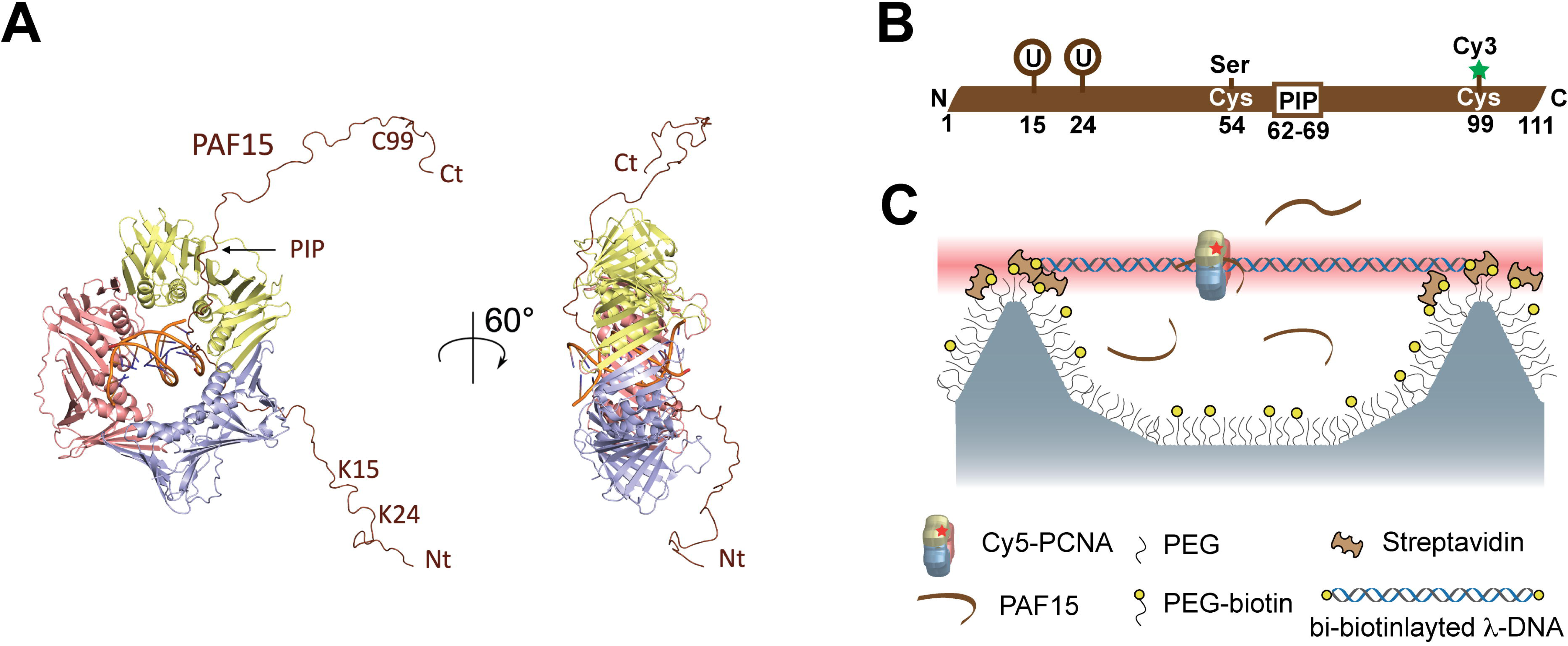
Structural model of ubiquitinated PAF15–PCNA–DNA complex and experimental schematic. **(A)** Cartoon representation of one PAF15 molecule bound to one protomer of the PCNA ring. The image is a composition of the crystal structure of PCNA bound to a primed DNA duplex and a PAF15 peptide (PDB: 6EHT) and a model of PAF15 bound to the PCNA ring based on NMR and SAXS measurements^18^. In the Di-UbPAF15 construct, residues 19 and 24 are covalently linked to one ubiquitin molecule each. **(B)** Schematic of the PAF15 sequence with Cy3 conjugated to the cysteine at position 99^th^ via a maleimide-thiol reaction. **(C)** Schematic of the experimental setup for detecting PCNA diffusion using DNA skybridge. A prism-type total internal reflection fluorescence microscopy generates a thin light sheet across the top ridges of the DNA skybridge, where DNA and PCNA molecules are localized.

Through its engagement with PCNA, PAF15 regulates the assembly and dynamic behavior of the replication machinery and coordinates the access of repair enzymes to damaged DNA sites. Still, the molecular mechanism of this regulation remains poorly understood. To address this gap, our study investigates the impact of PAF15 binding on PCNA’s diffusion and stability on DNA using single-molecule imaging techniques, such as the DNA skybridge^25^. By exploring how PAF15 influences PCNA’s interaction with DNA and modulates its availability for binding partners, we found that PAF15 binding significantly slows PCNA diffusion and stabilizes PCNA on DNA. Furthermore, we demonstrate that PAF15 directly modulates the ability of the RFC clamp loader to load PCNA onto DNA, by restricting the accessibility of the PCNA PIP-box binding sites. These effects suggest that PAF15 acts as a key regulatory factor capable of tuning PCNA’s functional state in response to cellular conditions. Furthermore, we propose PAF15’s role in modulating PCNA’s interactions with replication and TLS polymerases, influencing the balance between high-fidelity replication and lesion bypass. By elucidating the mechanisms underlying the PAF15-PCNA interaction, we provide new insights into how cells dynamically control PCNA’s function, with implications for understanding replication fidelity and the cellular response to DNA damage.

## Results

### PAF15 Slows PCNA Diffusion on DNA

To address the interaction of PAF15 with PCNA, we prepared Cy5-labeled PCNA and Cy3-labeled PAF15. A variant of PAF15 in which Cys54 was mutated to serine was site-specifically conjugated with a Cy3 fluorophore at Cys99 (**Fig. 1B**). Cy5-PCNA molecules were pre-loaded on λ-phage DNA molecules, which were biotinylated at both ends, stretched, and suspended within the DNA skybridge (**Figure 1C**). Upon introducing PAF15, we observed a significant slowdown in PCNA diffusion (**Figure 2A**). To quantify this effect, we determined the one-dimensional diffusion constant (*D*) of Cy5-PCNA by analyzing the mean square displacement (MSD) using MSD = 2*D*Δt, where Δt is the elapsed time, at various PAF15 concentrations. The results demonstrate that the PCNA diffusion constant decreases with increasing PAF15 concentrations by up to 15-fold (**Figure 2A**). This slowdown in diffusion cannot be solely attributed to the increased hydrodynamic size of the PCNA-PAF15 complex. A PCNA-Qdot complex, which has a much larger hydrodynamic size than the PCNA-PAF15 complex, only exhibited a ∼ 2-fold reduction in the diffusion constant^26^, suggesting an additional mechanism underlying the pronounced diffusion slowdown of PCNA in the presence of PAF15.

**Figure 2.**
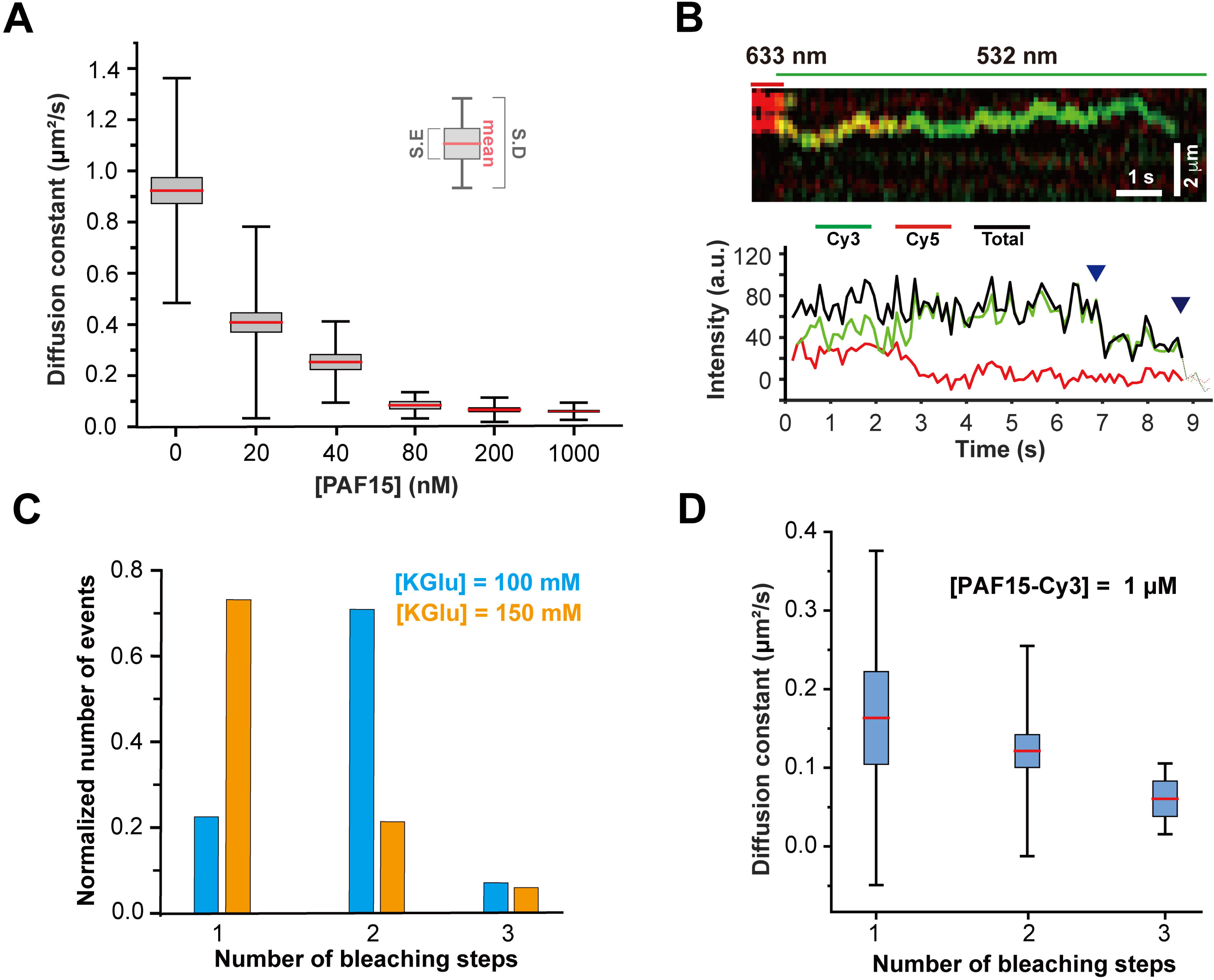
PAF15 modulates PCNA diffusion on DNA skybridge. **(A)** Box plot of PCNA diffusion constants measured at increasing concentrations of PAF15. Red lines indicate mean values, gray boxes represent standard error of the mean, and black whiskers denote standard deviation. Diffusion constants (mean ± s.e.m.) and sample sizes at various concentrations of PAF15 were as follows: points: 0.92 ± 0.05 μm^2^/s (n = 76) at 0 nM, 0.41 ± 0.04 μm^2^/s (n = 101) at 20 nM, 0.25 ± 0.03 μm^2^/s (n = 29) at 40 nM, 0.08 ± 0.01 μm^2^/s (n = 13) at 80 nM, 0.07 ± 0.01 μm^2^/s (n = 37) at 200 nM, and 0.061 ± 0.003 μm^2^/s (n = 119) at 1 μM. **(B)** Representative kymogram and fluorescence intensity trace showing two photobleaching steps in the presence of 1 μM PAF15-Cy3. In the kymogram, red indicates PCNA-Cy5 and green indicates PAF15-Cy3. In the intensity trace, green, red, and black lines correspond to Cy3, Cy5, and total intensity, respectively. **(C)** Histogram showing distributions of Cy3 photobleaching steps for Cy5-PCNA/PAF15-Cy3 complex under different potassium glutamate (KGlu) conditions with 1 μM PAF15-Cy3: 100 mM KGlu (cerulean) and 150 mM KGlu (light purple). Step distributions: 1 step 0.22 (100 mM), 0.73 (150 mM); 2 steps: 0.71 (100 mM), 0.21 (150 mM); 3 steps: 0.07 (100 mM), 0.06 (150 mM); n = 58 (100 mM) and n = 52 (150 mM). **(D)** Diffusion constants for PCNA-PAF15 complexes grouped by the number of PAF15-Cy3 molecules per complex: 0.16 ± 0.06 μm²/s (1 step, n = 13), 0.12 ± 0.06 μm²/s (2 steps, n = 41), 0.06 ± 0.02 μm²/s (3 steps, n = 4), and 0.13 ± 0.02 μm²/s (overall, n = 58).

Cy3-labeled PAF15 allowed us to monitor Cy3-photobleaching steps of PAF15-Cy3 colocalized with Cy5-PCNA (**Figures 2B and C**) and determine the diffusion constant of the complex, which varies based on the number of PAF15-Cy3 molecules bound to Cy5-PCNA (**Figure 2D**). The FRET signal between Cy3 and Cy5 confirmed the formation of the Cy5-PCNA/PAF-Cy3 complex, with two-step photobleaching indicating the binding of two PAF15-Cy3 molecules to Cy5-PCNA (**Figure 2B**). At a PAF15 concentration of 1 μM, where the PCNA diffusion constant reaches saturation, PAF15 predominantly associates with PCNA on DNA with a stoichiometry of one or two PAF15 molecules per ring, with rare instances of three PAF15 molecules bound to the same PCNA ring (**Figure 2C**). This depends on the ionic strength, as at a higher concentration of potassium glutamate (KGlu), the predominant stoichiometry is one PAF15 molecule per PCNA ring (**Figure 2C**). Only 6 ∼7 % of the PAF15 binding events are independent of the ionic strength. As more PAF15-Cy3 molecules bind to PCNA, the diffusion constant of the complex further decreases (**Figure 2D and Supplementary Figure S1**). Notably, the complex with two PAF15-Cy3 molecules exhibits around 7-fold slower diffusion (*D* = 0.13 ± 0.02 μm^2^/s) than PCNA alone (*D* = 0.92 ± 0.05 μm^2^/s) in the presence of 100 mM KGlu (**Figures A and 2D**). These findings demonstrate that the diffusion slowdown of PCNA is directly linked to PAF15 binding beyond simple hydrodynamic effects.

To further explore the impact of the PAF15-PCNA interaction on DNA sliding, we used three distinct PAF15 constructs: ΔN-PAF15, PAF15^50–77^, and Di-UbPAF15. The ΔN-PAF15 deletion mutant lacks the N-terminal 31 residues, the region that binds DNA and undergoes ubiquitination^10^. This construct allows us to investigate how PAF15’s DNA-binding ability influences PCNA diffusion. PAF15^50–77^ is a 28-residue-long peptide containing the PIP motif at the central region of PAF15. It binds to each of the three PIP-binding sites on the front side of the PCNA ring, but its N-terminal region is too short to come out of the ring^10^. The third variant, Di-UbPAF15, is doubly monoubiquitinated at Lys15 and Lys24. This form of PAF15 is predominantly found in the chromatin-associated fraction of cell extracts, but not in the soluble fraction^8^.

Di-UbPAF15 binds PCNA with a very similar affinity as PAF15 (K_d_ values are 1.6 and 1.1 μM at 298 K, respectively)^18,21^, while ΔN-PAF15 and PAF15^50–77^ bind with a 3-fold or a 5-fold lower affinity, respectively^18^. Therefore, we increased the concentration of each construct until the diffusion constant of Cy5-PCNA did not further decrease **(Supplementary Figure S2)**. Each PAF15 variant exhibited unique saturation concentration and diffusion constants **(Supplementary Figure S2)**. The saturation concentrations of PAF15 and ΔN-PAF15 differ more than expected from their similar PCNA-binding affinity, suggesting that the N-terminal disordered region of PAF15 plays a role in stabilizing the PCNA-PAF15 complex on DNA. At diffusion-saturation concentrations, the diffusion constants of the PCNA complexes followed this order: PAF15^50–77^ ≍ DiUb-PAF15 > ΔN-PAF15 > PAF15 **(Figure 3A)**. These results offer two key insights: (1) the N-terminal DNA binding region of PAF15 contributes to PCNA diffusion speed reduction, (2) the threading of the PAF15 chain inside the PCNA ring channel contributes the most to diffusion slowdown. Because the short PAF15^50–77^ does not completely thread through the ring and the ubiquitin moieties of DiUb-PAF15 cannot go through the gap between PCNA and DNA, we conclude that the N-terminal insertion and emergence on the back side of the ring must be the major factor in slowing down diffusion. These findings suggest that the association of PAF15 to DNA-loaded PCNA likely involves initial PIP-box binding followed by N-terminal insertion into the PCNA-DNA gap.

**Figure 3.**
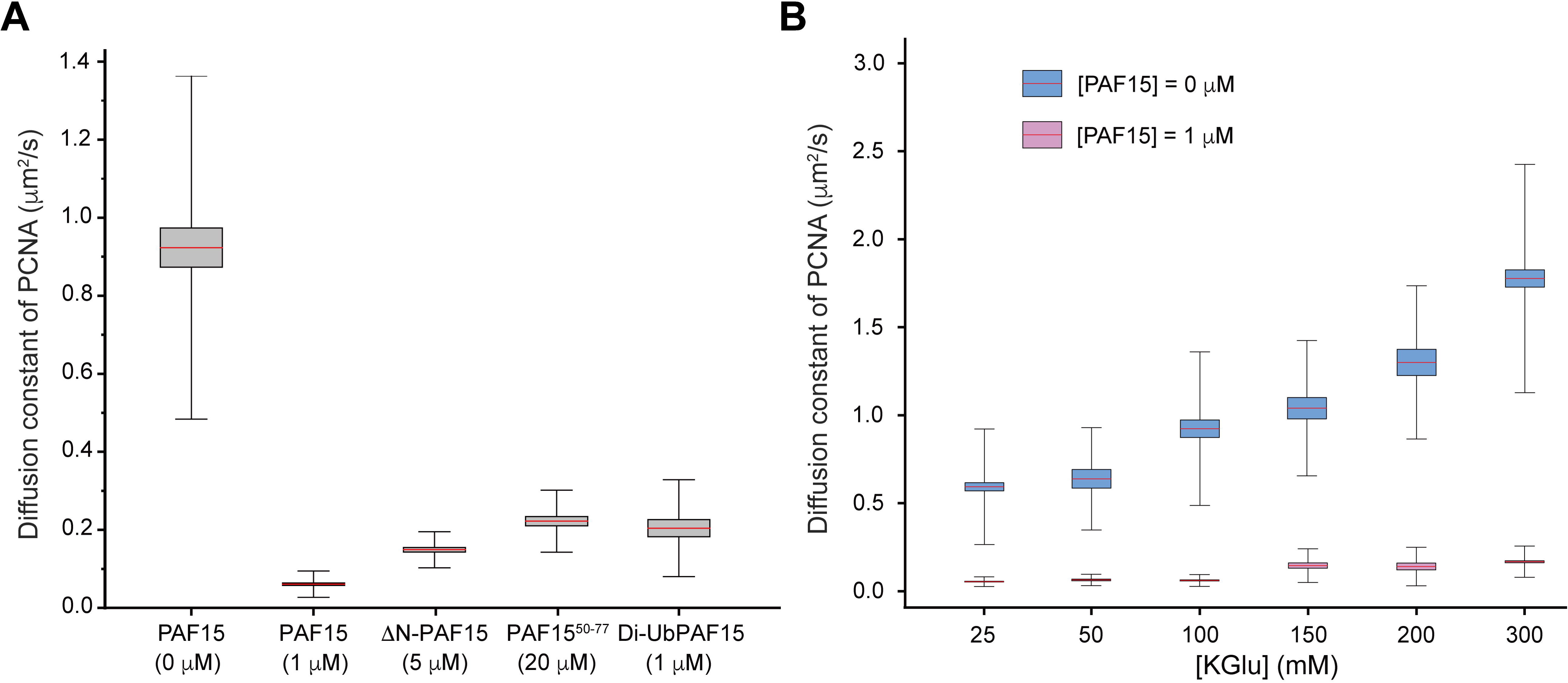
PAF15 constructs and ionic strength modulate PCNA diffusion. **(A)** Box plot showing the distribution of Cy5-PCNA diffusion constants with various PAF15 constructs and concentrations. Red lines indicate mean values, gray boxes and black whiskers represent the standard error and standard deviation, respectively. Diffusion constants (mean ± s.e.m.) and sample sizes (n) are as follows: 0.92 ± 0.05 μm^2^/s (no PAF15, n = 76), 0.061 ± 0.003 μm^2^/s (1 μM PAF15, n = 119), 0.15 ± 0.01 μm^2^/s (1 μM ΔN-PAF15, n = 61), 0.22 ± 0.01 μm^2^/s (20 μM PAF15/50-77, n = 40), 0.20 ± 0.02 μm^2^/s (1 μM Di-UbPAF15, n = 32). **(B)** Box plot showing Cy5-PCNA diffusion constants at varying KGlu concentrations. Diffusion constants without PAF15 and sample sizes are: 0.59 ± 0.02 μm^2^/s (n = 205) at 25 mM, 0.64 ± 0.05 μm^2^/s (n = 30) at 50 mM, 0.92 ± 0.05 μm^2^/s (n = 76) at 100 mM, 1.03 ± 0.06 μm^2^/s (n = 40) at 150 mM, 1.30 ± 0.07 μm^2^/s (n = 34) at 200 mM, and 1.78 ± 0.05 μm^2^/s (n = 174) at 300 mM. Diffusion constants with 1 μM PAF15: 0.054 ± 0.002 μm^2^/s (n = 205) at 25 mM, 0.063 ± 0.005 μm^2^/s (n = 43) at 50 mM, 0.061 ± 0.003 μm^2^/s (n = 119) at 100 mM, 0.145 ± 0.015 μm^2^/s (n = 40) at 150 mM, 0.140 ± 0.019 μm^2^/s (n = 32) at 200 mM, and 0.167 ± 0.006 μm^2^/s (n = 244) at 300 mM.

### PAF15 suppresses the rotation-uncoupled diffusion mode of PCNA on DNA

PCNA exhibits two distinct diffusion modes on the DNA duplex: rotation-coupled and rotation-uncoupled diffusion^27^. Rotation-coupled diffusion is defined as PCNA’s sliding by rotationally matching the pitch of the DNA helix, with basic residues lining the inner side of the ring establishing transient electrostatic interactions with the DNA phosphate groups^28^. Rotation-uncoupled diffusion refers to short periods of faster sliding, where those interactions are broken or weakened by water molecules, and the ring slides faster. Because the diffusion constants corresponding to each of these two modes have different dependencies on the ionic strength, their presence in the global diffusion can be experimentally detected. For this purpose, we conducted measurements at KGlu concentrations from 25 mM to 300 mM. In the absence of PAF15, the diffusion constant of Cy5-PCNA rings exhibited a gradual increase with KGlu concentration, confirming the occurrence of rotation-uncoupled diffusion (**Figure 3B**). In the presence of 1 μM PAF15, the diffusion constant remained the same from 25 to 100 mM KGlu and between 150 and 300 mM (**Figure 3B**), suggesting that PAF15 suppresses PCNA’s rotation-uncoupled diffusion. The diffusion constant significantly increased from 100 mM to 150 mM of KGlu concentration; we hypothesize that this increase is due to changes in the number of PAF15 molecules bound to PCNA (**Figure 2C**). These observations suggest that the PCNA’s rotation-uncoupled diffusion is suppressed regardless of the number of PAF15 molecules in the PAF15-PCNA complex.

To further support this interpretation, we quantitatively characterized the diffusion states of PCNA and PAF15-PCNA using the Bayesian inference framework combined with the theory of a multi-state Brownian motion process^29^. This analysis identifies the most probable diffusion model, including the number of diffusion states and their corresponding diffusion constants. We employed it to analyze the trajectory data of Cy5-PCNA, free or bound PAF15, to uncover their underlying diffusion states. Our hidden Markov model analysis confirmed that PCNA displays two distinct diffusion states, whereas PAF15-bound PCNA adopts a single diffusion state (**Supplementary Figure S3**). The PAF15’s unique ability to suppress rotation-uncoupled diffusion emphasizes its potential regulatory function along the DNA.

### Stabilization of DNA-loaded PCNA by PAF15 Binding

In our experimental setup, the Cy5-PCNA ring is loaded on the immobilized DNA by RFC, but PCNA can spontaneously unload by transient ring opening or protomer dissociation^30^. We measured the lifetime of PCNA on DNA to assess the effect of PAF15 binding on the stability of the assembly. For this purpose, we used the ratio of PCNA to bi-tethered DNA as an indicator of PCNA loading. This metric accurately reflects the amount of PCNA loaded on the DNA as it excludes PCNA unloading due to DNA detachment from the skybridge platform, which can occur over long times^31^.

To determine the amount of loaded PCNA on DNA, Cy5-PCNA, either alone or with PAF15, was first incubated with immobilized DNA in the presence of RFC and then washed to remove RFC and unloaded PCNA molecules. PCNA and Sytox Orange-stained DNA were imaged (**Supplementary Figure S4)** every two or five minutes, and the number of PCNA molecules on the DNA was counted. The resulting half-life time of PCNA on bi-tethered long DNA increased by more than 10-fold with the addition of PAF15 **(Figure 4; Methods)**. Notably, on average, the number of DNA molecules per image was reduced from 220 to 100 in 60 minutes after washing the RFC out. The amount of PCNA retained on the DNA skybridge in the presence of PAF15 may be underestimated due to breakage of the DNA. These results indicate that PAF15 substantially stabilizes the PCNA trimer on DNA, potentially contributing to processivity in DNA-associated processes.

**Figure 4.**
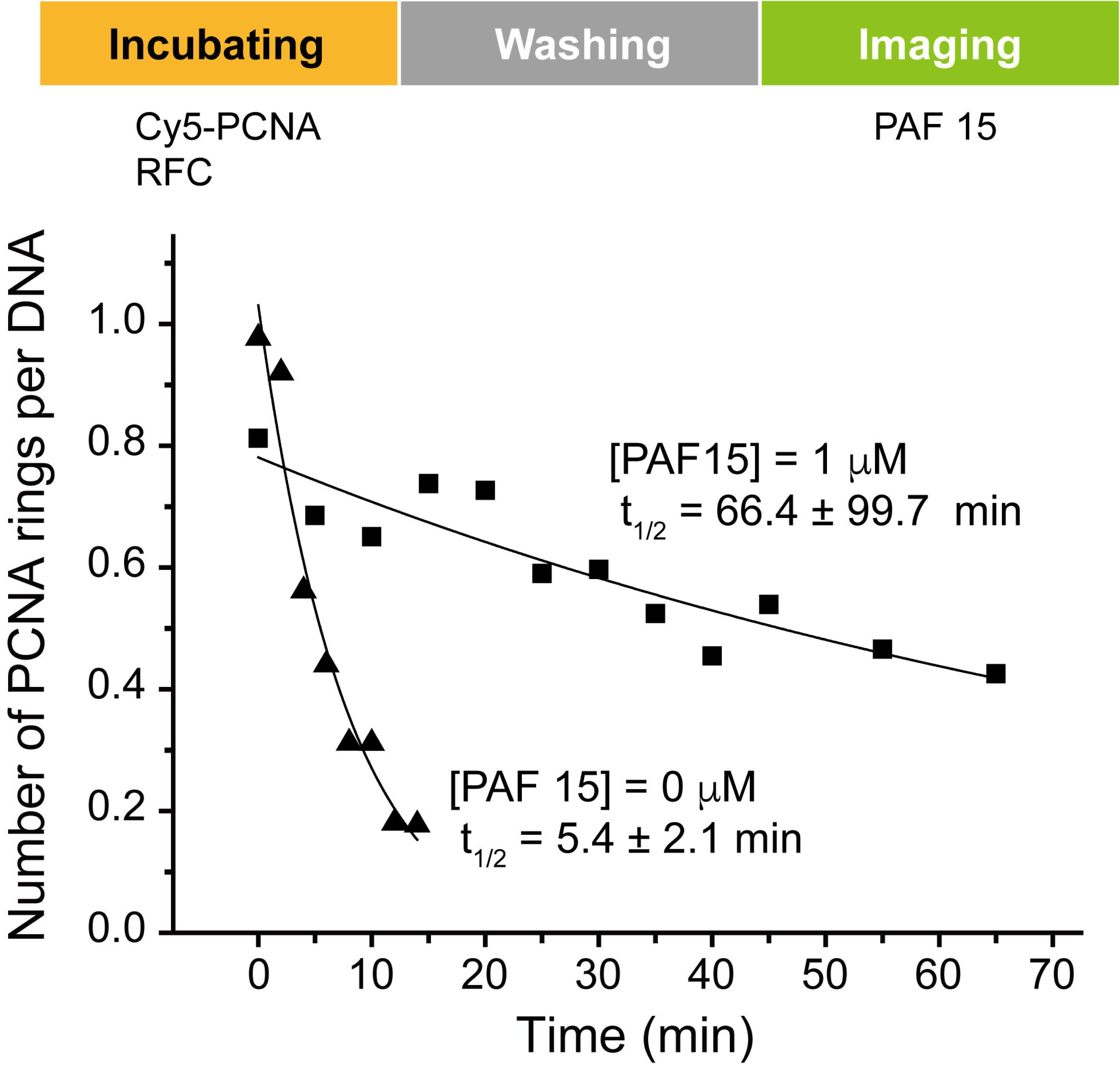
PAF15 decelerates spontaneous PCNA dissociation from DNA. The x-axis represents time, measured from the acquisition of the first image. Cy5-PCNA and Sytox Orange-stained DNA were imaged separately, and the number of PCNA and DNA was quantified over time. The resulting data were fitted to a first-order exponential decay model to determine the half-life of PCNA on DNA. Black triangles represent the condition without PAF15, and red squares represent the condition with 1 μM PAF15.

## Regulation of PCNA loading by PAF15

PCNA interacts with proteins containing PIP or PIP-like motifs, and the binding of PAF15 can compete with them. This is particularly relevant for protein complexes that involve multiple PIP interactions, such as the RFC complex, where the RFC1 and RFC3 subunits interact with two of the three PIP-binding sites of the PCNA ring ^32–34^. We hypothesized that PAF15 binding could affect RFC-mediated functions. To test this, we examined how varying concentrations of PAF15 influence PCNA loading. The amount of loaded PCNA was measured as the ratio of Cy5-PCNA to bi-tethered DNA. The results showed that high concentrations (> 80 nM) of PAF15 decreased PCNA loading, whereas an increase was observed at 40 nM **(Figure 5A)**. This concentration-dependent biphasic effect prompted us to investigate the unloading process as well.

**Figure 5.**
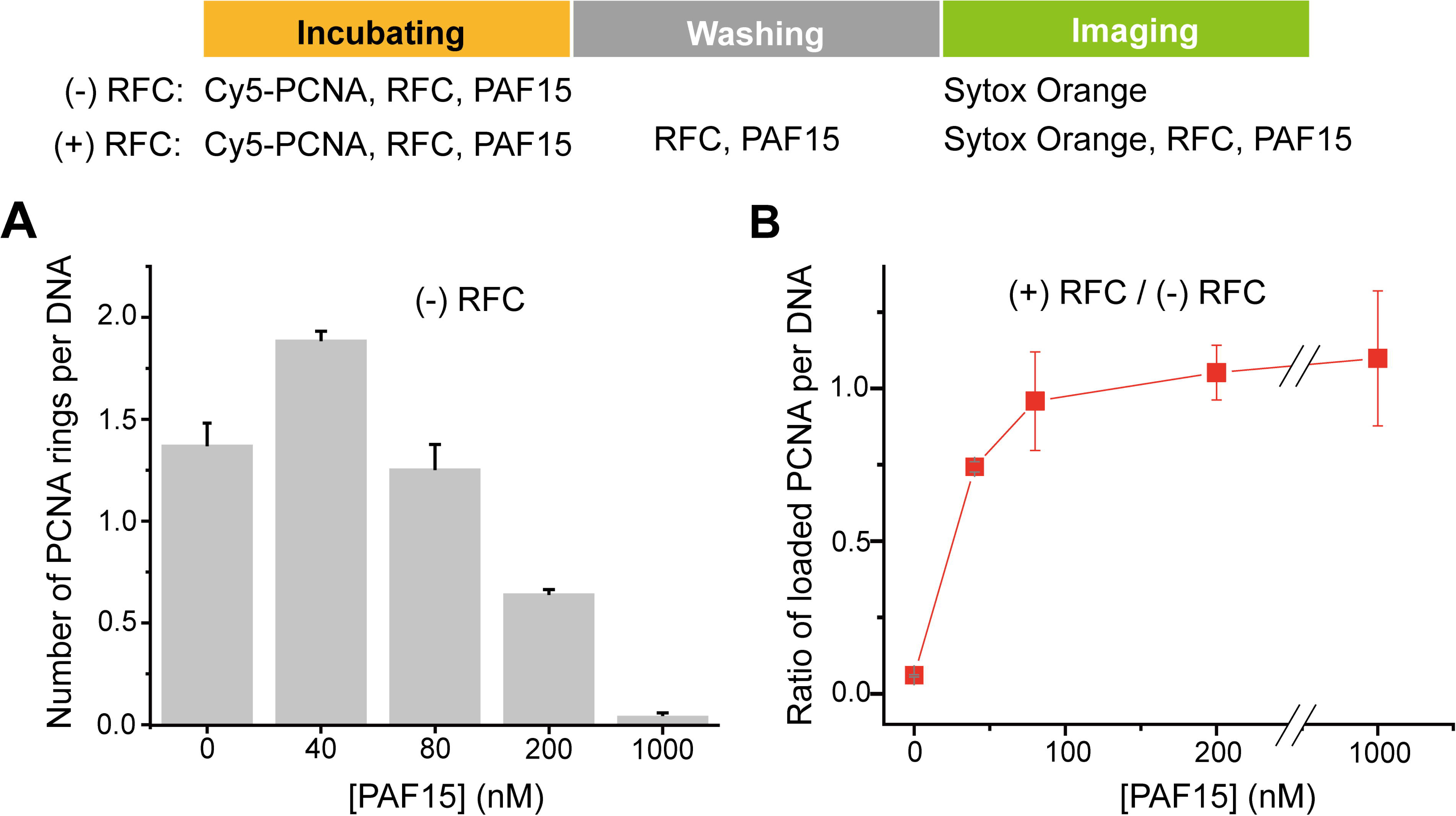
RFC and PAF15 regulate PCNA retention on DNA. The schematic illustrates the experimental procedure for two conditions: (–) RFC and (+) RFC. In both conditions, the incubation step includes PCNA, RFC, and the indicated concentration of PAF15. However, only the (+) RFC condition includes RFC and PAF15 in the washing and imaging steps. Data were acquired between 7 and 10 min after the initiation of the washing step. **(A)** Quantification of PCNA per DNA under the (–) RFC condition at varying PAF15 concentrations. Values represent mean ± s.e.m. and the number of independent experiments (N): 1.38 ± 0.11 (N = 3) at 0 nM, 1.89 ± 0.04 (N = 3) at 40 nM, 1.26 ± 0.12 (N = 4) at 80 nM, 0.65 ± 0.02 (N = 3) at 200 nM, and 0.045 ± 0.005 (N = 3) at 1000 nM. **(B)** Ratio of PCNA per DNA between the +(RFC, PAF15) and –(RFC, PAF15) conditions at the corresponding PAF15 concentrations. 0.061 ± 0.005 (N = 3) at 0 nM, 0.74 ± 0.02 (N = 3) at 40 nM, 0.96 ± 0.16 (N = 4) at 80 nM, 1.05 ± 0.09 (N = 3) at 200 nM, and 1.10 ± 0.22 (N = 3) at 1000 nM. The average number of bi-tethered DNA molecules per trial was approximately 220.

In a separate experiment, RFC and PAF15 were added to the three steps (incubating, washing, and imaging; **Figure 5B**). During the incubation step, RFC can both load and unload Cy5-PCNA^30^, but during the washing and imaging steps, it can only facilitate unloading, due to the absence of free PCNA. By comparing the amount of DNA-loaded PCNA in the two experiments (with or without RFC in the washing and imaging steps), we determined the RFC-mediated PCNA unloading/loading ratio at various PAF15 concentrations. The result is that RFC unloads PCNA rings from DNA, as expected, but PAF15 binding strongly reduces RFC-induced unloading, even at the lowest tested concentration **(Figure 5B)**.

During the incubation step, Cy5-PCNA loading and unloading reach a steady state over time. Although PAF15 inhibits both processes, it does so with differing efficiencies depending on its concentration. The diffusion rate of the PAF15-PCNA complex remains nearly constant at PAF15 concentrations above 80 nM **(Figure 2A)**, suggesting that PCNA is saturated with PAF15 at these concentrations. Since the affinity of PAF15 for free PCNA is not that high (K_d_ = 1.1 μM at 298 K)^18^, this finding indicates that the affinity of PAF15 for the DNA-loaded PCNA assembly is much higher than for free PCNA, implying that DNA enhances the overall binding affinity. This is consistent with the measured micromolar affinity of PAF15 for bare DNA^18^. Therefore, PAF15 at concentrations exceeding 80 nM effectively blocks PCNA loading by outcompeting 0.5 nM RFC during the incubation step. However, at lower concentrations (40 nM), PAF15 is sufficient to hinder unloading but not loading by RFC, shifting the steady state toward increased PCNA loading.

The increase of DNA-loaded PCNA lifetime caused by PAF15 **(Figure 4)** could cause an apparent increase in loading at 40 nM PAF15 in the experiment without RFC and PAF15 in the washing and imaging solution **(Figure 5A)**. To clarify if that is the case, we conducted a complementary experiment by adding PAF15 during the washing and imaging steps (**Supplementary Figure S5**). This approach minimized lifetime differences, yet the results remained consistent with **Figure 5**, confirming that the larger binding affinity of PAF15 to DNA-loaded PCNA compared to free PCNA is key to the enhanced loading.

We also measured the RFC-mediated unloading rate of Cy5-PCNA from DNA under our experimental conditions. In the absence of PAF15, most loaded PCNA dissociated from DNA within seven minutes (**Supplementary Figure S6**), which is consistent with the data in **Figure 5B**. Finally, we confirmed that any nonspecific interactions between free PAF15 and RFC had minimal impact on the PCNA unloading experiment. We incubated DNA-loaded Cy5-PCNA with PAF15, washed away the unbound PAF15, and then monitored PCNA dissociation in the presence of RFC (**Supplementary Figure S6**). The results indicated that PAF15 bound to DNA-loaded PCNA effectively blocked RFC-mediated unloading for an extended period. These results indicate that PAF15 binding to PCNA can modulate PCNA loading and unloading, balancing these processes based on PAF15 concentration and influencing PCNA availability on DNA.

## Discussion

The results reveal that PAF15 slows down PCNA diffusion on the DNA up to 15-fold in a manner dependent on the PIP-box binding and on the threading of its N-terminal region in the hole of the PCNA ring. The insertion of this region in the PCNA-DNA gap acts like a wedge, increasing the friction that opposes free diffusion. Pioneering single-molecule experiments showed that the stochastic displacement of PCNA along the DNA is predominantly coupled with ring rotation, slowly tracking the helical pitch of the duplex, while less frequently, PCNA fast moves uncoupled with rotation^27^. While coarse-grained computational simulations^35^ suggest that the rotation-uncoupled translation is predominant, both experimental and computational simulations support a predominant rotation-coupled translation driven by transient electrostatic interactions between positively charged side chains lining the inner surface of the ring and the phosphate groups of the duplex^28,36^. Our results are consistent with previous experimental measurements and show that the friction caused by PAF15 insertion suppresses the faster rotation-uncoupled translation. The diffusion slowdown may be critical in processes where PCNA’s movement impacts the outcome of enzyme reactions, such as DNA replication and repair. Although PAF15-bound PCNA diffusion remains approximately 15-fold faster than the DNA synthesis rate by polymerase δ (∼ 1,500 bp/s vs. ∼ 100 bp/s)^37^, the ∼7-fold reduction in diffusion caused by PAF15 binding significantly narrows this gap. This reduction may functionally influence lagging-strand synthesis by bringing the rate of PCNA translocation closer to the pace of DNA synthesis, potentially affecting the coordination of polymerase progression and Okazaki fragment processing.

We found that PAF15 binding increases the lifetime of DNA-loaded PCNA. This stabilization is consistent with recent cellular assays showing that PAF15 depletion reduced the residence time of PCNA in the chromatin while ectopic expression stabilized it^15^. Our results also demonstrate that PAF15 can modulate RFC-mediated PCNA loading and unloading in a concentration-dependent manner. This finding suggests that PAF15 cellular concentration could dynamically control PCNA availability on DNA, with high levels favoring stable PCNA binding and low levels permitting RFC-mediated turnover. PAF15 is about 7-fold less abundant than PCNA inside the nuclei of human cells^15^, but its concentration is regulated through the cell cycle, peaking in the S phase (and mostly chromatin-bound) and declining after G2 phase (during which it is released into the nuclear soluble fraction)^26^. Such regulation could be important for orchestrating DNA-loaded PCNA availability during different stages of DNA processing. Ectopically increasing PAF15 to very high levels leads to a lethal replication response in human cells, and PAF15 is found overexpressed in most tumor types^15^.

The interaction between PAF15 and PCNA is likely tailored to meet the specific demands of DNA synthesis and repair. PAF15 could occupy and thereby restrict one of PCNA’s PIP-box motifs, potentially limiting the recruitment of other PIP-box-containing proteins. For instance, polymerase ε requires all three PIP-boxes of PCNA for optimal interaction, and blocking one or two of them would prevent the functional polymerase ε-PCNA interaction^38^. The presence of PAF15 may thus play a role in selectively modulating PCNA’s interactions with its many partners, fine-tuning PCNA’s activity in a context-dependent manner. Recent experiments suggest that PAF15 is a low-dosage factor critical for lagging-strand replication, and not for leading-strand replication. The DNA lagging strand is replicated as short Okazaki fragments by DNA polymerase δ, with a PIP motif bound to one of the three PIP-binding sites of the PCNA ring^9^. The maturation of the Okazaki fragments into a continuous strand requires the action of Flap endonuclease 1 (FEN 1) and Ligase 1 (Lig 1), which also bind to PCNA through their PIP motifs^9,10^. The so-called toolbelt model proposes that PCNA simultaneously binds more than one of those enzymes, facilitating the handing off of the DNA from one to another, and PCNA-bound PAF15 affects how this handing off occurs^8^. Most of the chromatin-bound PAF15 is monoubiquitinated at Lys15 and Lys24^24^. PAF15 ubiquitination recruits DNA methyl transferase 1 to the replication fork to restore the methylation pattern in the newly synthesized strand^20^ and triggers the switch between replicative to TLS polymerases at DNA damage sites by its degradation. Our experiments cannot provide conclusive evidence for the possible impact of this posttranslational modification on PCNA diffusion.

Overall, our study reveals that PAF15 regulates PCNA’s diffusion and stability on DNA, providing new insights into the molecular mechanism underlying its impact on DNA replication and repair in human cells. Future research should investigate the potential therapeutic implications of targeting PAF15-PCNA interactions, especially in conditions where TLS or replicative polymerases are aberrantly regulated, to maintain genome stability.

## Materials and methods

### RFC preparation

Human full-length replication factor C (RFC) expression vectors, pGBM-RFC1, pET-RFC4/2, and pCDFK-RFC5/3, were generous gifts from Dr. Yuji Masuda^39^. To generate an N-terminus truncated version of RFC1, the coding region for the first 550 amino acids was deleted and replaced with a Strep II–tag coding sequence, resulting in the construct pGBM-ΔN-RFC1-Strep II. Additionally, pCDFK-RFC5/3 was modified to encode a 6xHis tag at the N-terminus of RFC3. These three plasmids were co-transformed into *E. coli* BL21(DE3) cells, and transformants were selected on LB agar plates containing kanamycin, ampicillin, and streptomycin.

RFC was overexpressed by culturing transformed cells in 8 L TB supplemented with the same three antibiotics. Cells were grown at 25 °C to an OD_600_ of 0.8, then induced with 0.2 mM IPTG and incubated for an additional 24 h at 16 °C. Cells were harvested by centrifugation and resuspended in Buffer A supplemented with 600 mM NaCl (Buffer A: 50 mM HEPES-KOH, pH 7.5, 10 mM β-mercaptoethanol, and 1 mM PMSF). All subsequent steps were performed at 4 °C.

Cell lysis was carried out enzymatically using 2 mg/mL lysozyme and mechanically by sonication. The lysate was clarified by centrifugation at 35,000 rpm for 50 min. The supernatant was adjusted to 10 mM imidazole and applied to a 5 mL HisTrap HP column (GE Healthcare). Elution was carried out using a linear gradient from Buffer A to Buffer A containing 300 mM imidazole. Fractions containing all RFC subunits were pooled and passed through a 1mL StrepTrap HP column (GE Healthcare), which retains complexes with incorrect stoichiometry; correctly assembled RFC complexes appeared in the flow-through.

The flow-through was diluted to 300 mM NaCl with Buffer A and loaded onto a 1 mL HiTrap Heparin HP column. Elution was performed using a linear gradient from Buffer A with 300 mM NaCl to Buffer A with 1 M NaCl. Fractions containing RFC subunits were collected, concentrated, and subjected to gel filtration using a HiLoad Superdex 16/600 200 column equilibrated in storage buffer (25 mM Tris–HCl, pH 8.0, 250 mM NaCl, 0.01% NP-40, 1 mM DTT, 0.5 mM EDTA, and 10% glycerol). Fractions containing RFC with correct stoichiometry co-eluted, and were collected, concentrated, and flash-frozen for storage.

### PCNA preparation

The PCNA open reading frame (ORF) was cloned into the MCS1 site of the pET-Duet1 vector (Novagen), resulting in the expression of PCNA with an N-terminal 6×His tag. PCNA was purified using a series of chromatographic steps, including HisTrap HP, Q-Sepharose, and Superdex 16/600 75 size-exclusion chromatography. Gel filtration was performed using buffer containing 50 mM HEPES–KOH (pH 7.5), 100 mM NaCl, and 0.5 mM TCEP. For fluorescent labeling, purified PCNA was incubated with a five-fold molar excess of Cy5 Maleimide (GE Healthcare) for 6 h at 4 °C. The reaction was quenched by adding DTT to a final concentration of 10 mM. Excess unreacted dye was removed by gel filtration using a Superdex 75 10/300 GL column. Fractions containing Cy5-labeled PCNA were pooled and dialyzed against storage buffer (25 mM Tris–HCl, pH 7.5, 50% glycerol, 50 mM NaCl, 1 mM EDTA, and 1 mM DTT).

### PAF15 preparation

Human PAF15 (UniProt: Q15004) and the truncated PAF15 variant, lacking residues 2-31 (ΔN-PAF15), were produced in *E. coli* BL21(DE3) cells and purified through three sequential chromatographic steps: anion exchange, reverse phase, and cation exchange chromatography, as previously detailed^18^. The Cys54Ser mutant clone was generated using the QuikChange Site-Directed Mutagenesis Kit (Agilent), and protein was produced and purified in the same way as the wild type. Mass spectrometry confirmed the identity of the three pure proteins, without extraneous residues but lacking the initial methionine. For the preparation of doubly monoubiquitinated PAF15, a mutant protein Lys15Cys, Lys24Cys, Cys54Ser, and Cys99Ser was obtained from a synthetic clone and disulfide-linked with ubiquitin (with a Cys residue after the C-terminal Gly76 activated with cysteamine) as previously described in detail^21^. Mass spectrometry confirmed the identity of the protein, which lacked the initial methionine.

### PAF15^50–77^ preparation

The PAF15^50–77^ peptide (sequence: GNPVCVRPTPKWQKGIGEFFRLSPKDSE) was purchased from Apeptide. Concentrated stock solutions were prepared by dissolving the lyophilized peptide in water, and the pH was adjusted to approximately 7.0 using NaOH. These stocks were diluted in buffered solutions as needed. Peptide concentrations were determined by measuring absorbance at 280 nm, using an extinction constant calculated from the amino acid composition.

### Single-molecule fluorescence imaging

DNA substrates (10 pM) prepared in blocking buffer (**Supplementary Table S1**) were infused into a flow chamber (30 mm length x 4 mm width x 0.10 mm height) at a flow rate of 0.06 ml/min using a syringe pump (Harvard Apparatus). To initiate loading, 1 nM Cy5-PCNA and 1 nM RFC, prepared in reaction buffer (**Supplementary Table S1**), were introduced into the chamber and incubated for 1 min to allow PCNA loading onto DNA molecules. Following incubation, unbound PCNA and RFC were removed by washing with 1x reaction buffer.

Imaging of Cy5-PCNA bound to DNA was performed in imaging buffer (**Supplementary Table S1**), supplemented with an oxygen scavenging system to minimize photobleaching and suppress photoblinking of the Cy5 fluorophore. Experimental conditions, such as potassium glutamate or PAF15 concentration in imaging buffer, were adjusted as needed.

Fluorescence emission from Cy5 was excited with a 638 nm DPSS laser (Cobolt Samba, 100 mW) and imaged through a 1.6x magnifier on a prism-type total internal reflection fluorescence microscope (Olympus IX-71) equipped with a 60x water-immersion objective (Numerical Aperture = 1.2). Images were captured with an EMCCD camera (ImagEM C9100-13, Hamamatsu) and recorded using MetaMorph 7.6 (Molecular Devices) at a 100-ms time resolution. DNA molecules stained with Sytox Orange (Thermo Fisher Scientific) were excited with a 532 nm DPSS laser (Cobolt Samba, 100 mW). Data analysis were performed using DiaTrack 3.04^40^ and MATLAB 2013b (MathWorks) with a radial symmetry algorithm^41^. All experiments were conducted at room temperature (approximately 22°C).

### Two-color imaging of PAF15-Cy3 bleaching step test

The microscopy setup and experimental procedures were identical to those used for single-molecule fluorescence imaging, up to the PCNA incubation step (see Methods). Following PCNA incubation, 1 μM PAF15-Cy3 in reaction buffer was introduced into the flow chamber and incubated for 10 min. Unbound PAF15-Cy3 was then removed by washing twice with 400 μL of reaction buffer. After washing, imaging buffer was introduced to the chamber, and imaging started.

Only images acquired between 10 and 15 min after the start of the washing step were analyzed to ensure consistent natural dissociation behavior. A minimum 10-min waiting period was required to complete the wash and allow flow stabilization. The potassium glutamate concentration used from PAF15-Cy3 incubation through imaging was either 100 mM or 150 mM, as specified in the corresponding data.

Co-localization of Cy3 and Cy5 fluorescence signals was determined using a custom MATLAB script [2013b (MathWorks)] based on the THIN-PLATE SPLINE (TPS) algorithm. Particle tracking was performed using Diatrack 3.04 and the radial symmetry algorithm. Fluorescence intensities were quantified around the tracked positions for each fluorophore.

### PCNA half-life time determination

To determine the half-life time of PCNA on DNA, we counted the number of PCNA molecules diffusing on DNA, and DNA molecules were counted over time. We modeled the dissociation process of PCNA from DNA as a first-order exponential decay, represented by 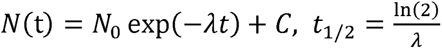, where *N(*t) is the number of PCNA molecules per DNA over time, *N*_0_ is the initial number of PCNA, *C* is the error term, is the decay constant, and *t*_1/2_ is the half-life time calculated.

### Measuring PCNA loading and unloading modulated by PAF15

For the experiment, an incubation solution containing 0.5 nM PCNA-Cy5, 0.5 nM RFC, and PAF15 at the indicated concentration, and imaging buffer was introduced into the flow chamber. After incubation, the solution was washed out using a condition-specific washing solution: in the (–) RFC condition, this consisted of reaction buffer alone; in the (+) RFC condition, the wash contained reaction buffer supplemented with 0.5 nM RFC and the indicated concentration of PAF15.

Following the wash, an imaging buffer was introduced for imaging. For (–) RFC condition, this buffer included imaging buffer and 1 nM Sytox Orange (Thermo Scientific). For the (+) RFC condition, the imaging buffer also included imaging buffer, 1 nM Sytox Orange, 0.5 nM RFC and the corresponding concentration of PAF15.

The incubation step lasted 3.5 min, followed by a 4.5 min wash and 1.5 min buffer exchange for imaging. To ensure consistent analysis of PCNA dissociation, image stacks were collected from 7 min to 10 min after the end of the incubation step. Two-color image stacks were acquired at the same field of view using separate excitation with 638 nm and 532 nm lasers to visualize PCNA-Cy5 and Sytox Orange-stained DNA, respectively. The number of fluorescent PCNA and DNA molecules within the imaging field was then quantified.

### Synthesis of bi-biotinylated λ DNA

Bi-biotinylated λ DNA was prepared by ligating two distinct oligonucleotides (**Supplementary Table S2**) to the 12-nt cohesive ends at both termini of λ-phage DNA. To prevent self-ligation between the oligos, two ligation steps were performed sequentially. In the first step, λ-phage DNA was mixed with the 3’biotin-#1 oligo at a 1:20 molar ratio in ligation buffer. In the second step, the 3’biotin-#2 oligo was added at a five-fold molar excess relative to the 3’biotin-#1 oligo, which resulted in a final ratio of λ-phage DNA:3’ biotin-#1:3’ biotin-#2 = 1:20:100. Following the second ligation, excess free oligos were removed using a Float-A-Lyzer® G2 (Spectra-Por), yielding purified bi-biotinylated λ-phage DNA suitable for DNA skybridge applications.

## Author contributions

D.K., F.J.B., A.D.B., and J.-B.L. conceptualized the study. D.K. and G.B. performed all single-molecule experiments. A.G.-M., A.D.B., and F.J.B. provided PAF15, fluorescently labeled PAF15, and PAF15 variants. M.T. and S.M.H. supplied RFC and PCNA. A.L.C. and J.-H.J conducted theoretical studies. C.J. provided the 3D-structured quartz for the DNA skybridge. All authors participated in data analyses. D.K., F.J.B., A.D.B., and J.-B.L. wrote the manuscript with input from all authors.

## Data availability

The data underlying this article are available upon request from the corresponding author(s).

