## Supplementary Information for "PAF15 stabilizes PCNA on DNA and slows down its sliding dynamics"

**Supplementary table S1.** Composition of the buffers used in the experiments.

| Name | Contents |
|---|---|
| Blocking buffer | 20 mM Tris-HCl, pH 7.5, 50 mM NaCl, 2 mM EDTA, 0.0025% Tween20 |
| Reaction buffer | 25 mM Tris-HCl, pH 7.4,100 mM KGlu, 5 mM $MgCl_2$, 0.0025% Tween20, 1 mM ATP, 0.1 mM ADP, 1 mM DTT, 0.1 mg/ml BSA |
| Imaging buffer | 25 mM Tris-HCl, pH 7.4,100 mM KGlu, 5 mM $MgCl_2$, 0.0025% Tween20, 1 mM ATP, 0.1 mM ADP, 1 mM DTT, 0.1 mg/ml BSA, 2 mM Trolox, 5 mM Protocatechuic acid (PCA), 200 nM Protocatechuate 3,4 dioxygenase (PCD) |

**Supplementary table S2.** Oligonucleotide sequences and modifications used for λ-phage DNA immobilization.

| **Bi-biotin λ–phage DNA** | |
|---|---|
| **Name** | **Sequence and modification** |
| 3’biotin–#1 | **5’**– /5Phos/AGG TCG CCG CCC TT/3Bio/ –**3’** |
| 3’biotin –#2 | **5’**– /5Phos/GGG CGG CGA CCT TT/3Bio/ –**3’** |

/3Bio/: 3’ biotin modification, /5Phos/: 5' Phosphorylation

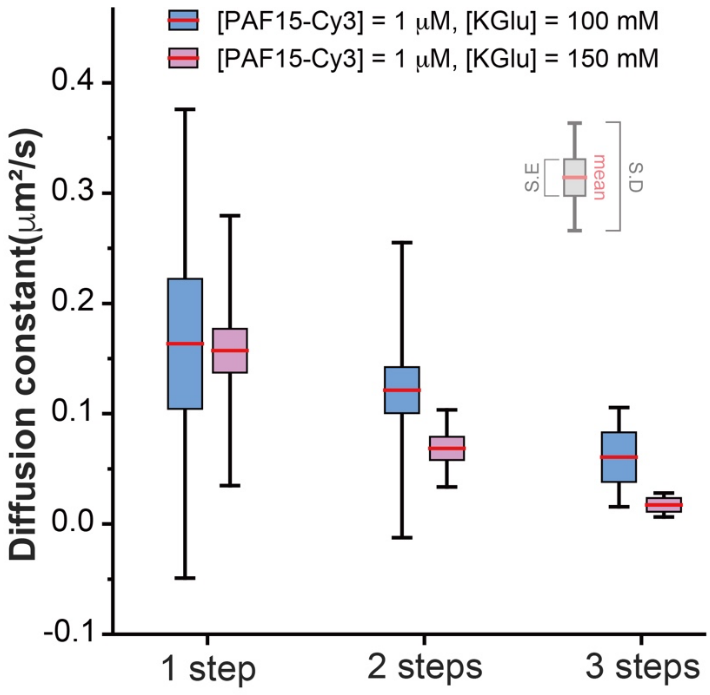


**Supplementary Figure S1**. **Diffusion constants as a function of bleaching step and salt concentration.** This plot shows the diffusion constants of the Cy5-PCNA and PAF15-Cy3 complex under different potassium glutamate (KGlu) concentrations and bleaching steps. The measured diffusion constants (mean ± s.e.m.) are as follows:

- 1 step: 0.16 ± 0.06 μm²/s (100 mM KGlu, *n* = 13), 0.16 ± 0.02 μm²/s (150 mM KGlu, n = 38)
- 2 steps: 0.12 ± 0.06 μm²/s (100 mM KGlu, *n* = 41), 0.069 ± 0.010 μm²/s (150 mM KGlu, n = 11)
- 3 steps: 0.061 ± 0.020 μm²/s (100 mM KGlu, *n* = 4), 0.017 ± 0.005 μm²/s (150 mM KGlu, n = 3)

Here, n denotes the number of PAF15-Cy3 molecules analyzed.

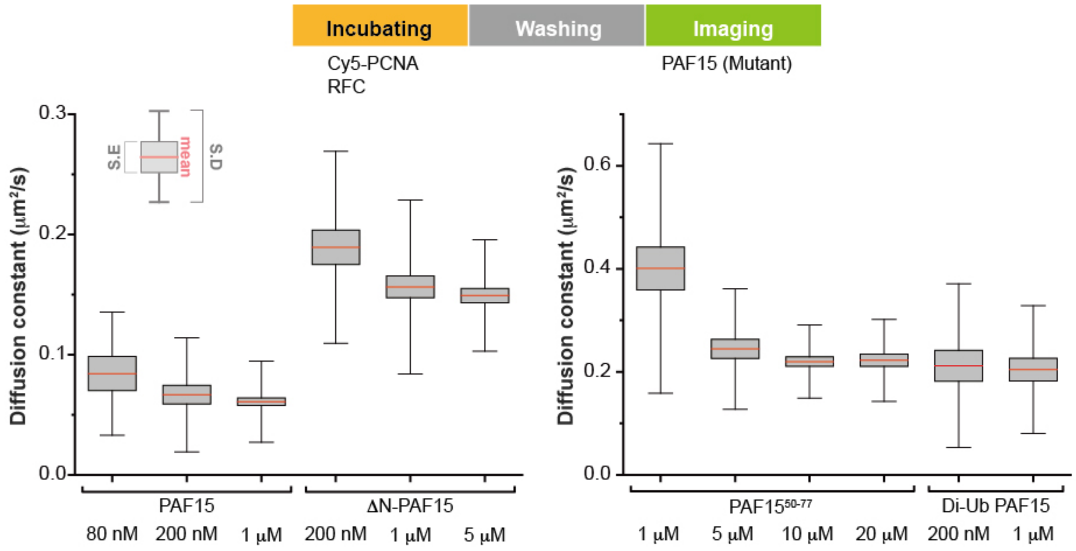


**Supplementary Figure S2. Effect of PAF15 constructs and their concentrations on Cy5-PCNA diffusion slowdown.** The diffusion constants were measured as increasing concentrations of each PAF15 variant were added, up to the point where no further significant reduction in PCNA diffusion was observed—interpreted as the saturation point for each variant. The diffusion constants under each condition, expressed as the mean ± standard error of the mean are as follows:

- **PAF15:**
  - 80 nM: 0.0843 ± 0.0038 μm²/s (n = 13), 200 nM: 0.0667 ± 0.0013 μm²/s (n = 37), 1 μM: 0.0610 ± 0.0003 μm²/s (n = 119).
- **ΔN-PAF15:**
  - 200 nM: 0.1893 ± 0.0025 μm²/s (n = 31), 1 μM: 0.1564 ± 0.0011 μm²/s (n = 64), 5 μM: 0.1493 ± 0.0008 μm²/s (n = 61).
- **PAF15$^{50–77}$ fragment:**
  - 1 μM: 0.4009 ± 0.0070 μm²/s (n = 34), 5 μM: 0.2447 ± 0.0029 μm²/s (n = 40), 10 μM: 0.2201 ± 0.0012 μm²/s (n = 57), 20 μM: 0.2225 ± 0.0017 μm²/s (n = 45).
- **Di-Ub PAF15:**
  - 200 nM: 0.2120 ± 0.0056 μm²/s (n = 28), 1 μM: 0.2044 ± 0.0038 μm²/s (n = 32).

Here, n denotes the number of Cy5-PCNA molecules analyzed.

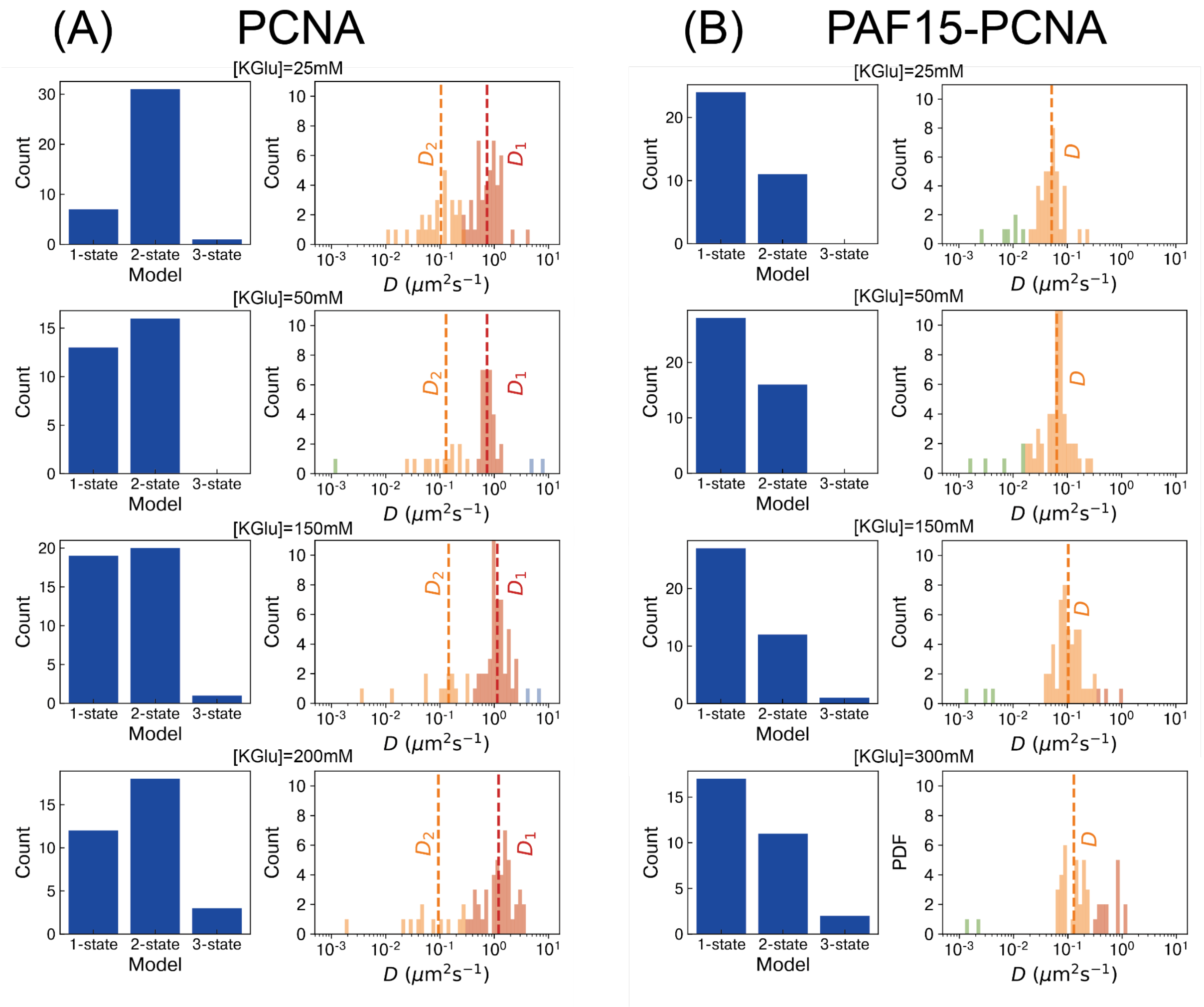


**Supplementary Figure S3: Diffusion state analysis of (A) PCNA and (B) PAF15-PCNA.** The left panels in (A) and (B) show the optimal model selection using the Akaike information criterion (AIC) method for PCNA and PAF15-PCNA at different salt concentrations of KGlu, as indicated at the top of each panel. In panel (A), the AIC consistently identifies a two-state diffusion model as the best fit model for PCNA dynamics across all KGlu concentrations. However, panel (B) shows that for PAF15-PCNA, the AIC reveals a one-state diffusion model as the optimal model across all KGlu concentrations. To further characterize the diffusion states of PCNA and PAF15-PCNA, we use a Bayesian Gaussian mixture model (BGMM) to identify the dominant modes of the Gaussian components that provide distinct diffusion coefficient values. The right panels in (A) and (B) present the clustering results of the estimated diffusion coefficients using the maximum likelihood estimator for PCNA and PAF15-PCNA at the given KGlu concentrations. In panel (A), the BGMM analysis reveals two distinct clusters (colored red and orange) of the estimated diffusion coefficients for PCNA. The modes of these clusters, indicated by dashed red and orange lines, correspond to the diffusion coefficients $D_1$ and $D_2$ of the two-state diffusion model. The diffusion coefficients $D_1$ and $D_2$ likely correspond to fast rotation-uncoupled translational diffusion and slow rotation-coupled translation, respectively. Panel (B) shows that the BGMM identifies a single dominant cluster (orange colored) of diffusion coefficients for PAF15-PCNA, with the major mode (indicated by orange dashed lines) corresponding to the diffusion coefficient $D$. At the lowest KGlu concentration (25 mM), we also observe a minor cluster (green colored) with a diffusion coefficient of ~ 0.01 $\mu m^2 s^{-1}$. Interestingly, the red clusters (associated with $D_1$) in the clustering plots of the PCNA-only are absent in the corresponding plots of PAF15-PCNA, except at the highest salt concentration. At [KGlu] = 300 mM, a red cluster with diffusion coefficient $\approx$ 0.49 $\mu m^2 s^{-1}$ appears, possibly implying a few trajectories with faster diffusion. Even in this case, the diffusion coefficient value is lower than the $D_1$ ($\approx$ 0.73 $\mu m^2 s^{-1}$) observed for PCNA alone at the lowest KGlu concentration (25 mM). These results indicate that PAF15 binding to PCNA suppresses the higher diffusion coefficient component ($D_1$), implying the suppression of the faster, rotation-uncoupled translation. Overall, our results demonstrate that a one-state model is the dominant diffusion model for PAF15-PCNA dynamics.

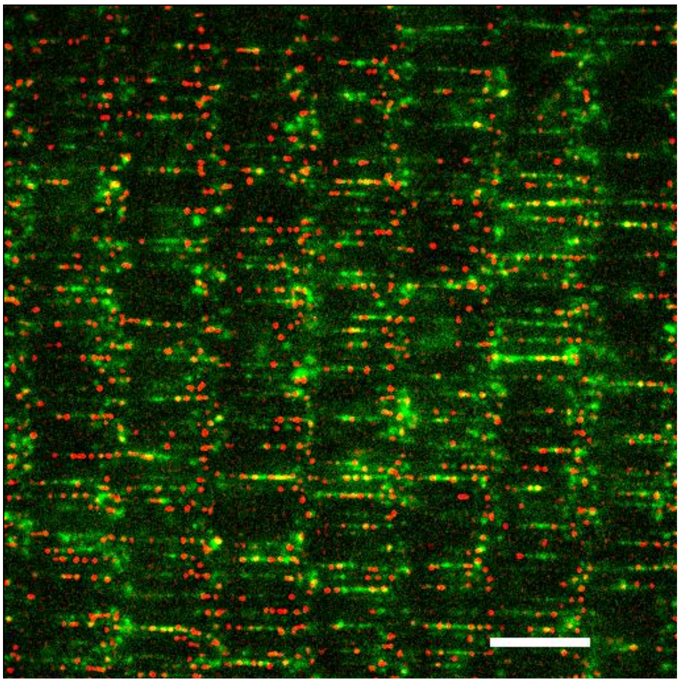

**Supplementary Figure S4. Representative image of the PCNA loading**. The red spots represent Cy5-PCNA, and the green line indicates DNA stained with Sytox Orange. This image is part of the dataset shown in Figure 5C, specifically representing the –RFC condition at [PAF15] = 40 nM. In this field of view, 511 PCNA molecules and 233 DNA molecules were detected. The white scale bar corresponds to 13 µm.

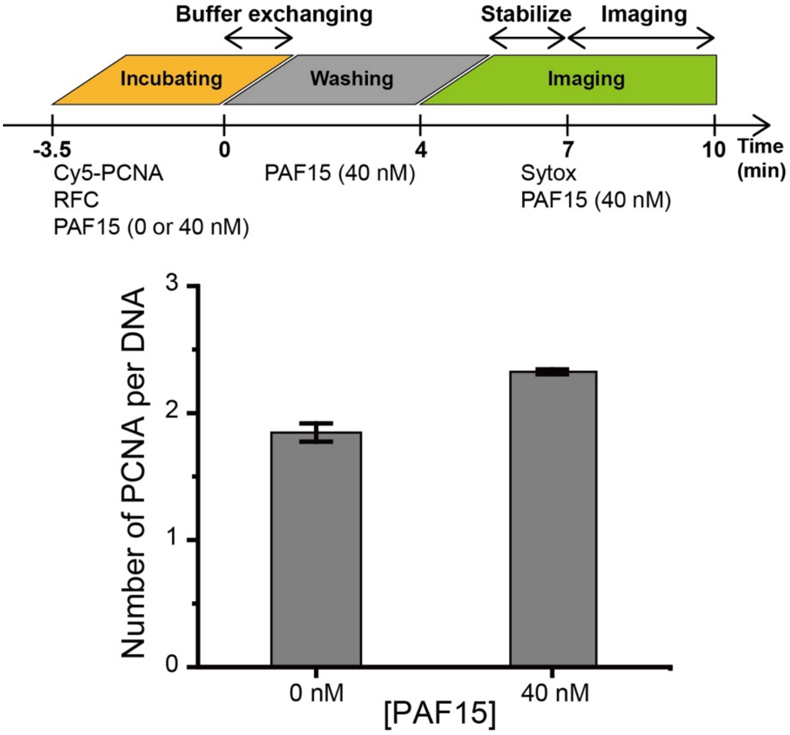


**Supplementary Figure S5. Comparison of PCNA loading levels in the absence or the presence of PAF15 at low concentration**. This experiment compares the amount of PCNA loaded per DNA molecule under two conditions: absence of PAF15 and low PAF15 concentration (40 nM), with both samples processed using the same washing and imaging steps. Figure 5a suggested that increased PCNA loading at low PAF15 concentration may result from differential binding affinities between free and DNA-loaded PCNA. However, the increased lifetime of PCNA observed in Figure 4 could cause an unrelated apparent increase. To test this, we performed an experiment similar to that shown in Figure 5a but adding PAF15 during the washing and imaging steps and controlling for potential imaging artifacts by keeping washing and imaging conditions identical. The number of PCNA rings per DNA (mean ± SD) is 1.846 ± 0.071 at 0 nM PAF15 (n = 3), 2.324 ± 0.020 at 40 nM ÂF15 (n = 3). Here, n denotes the number of independent experiments.

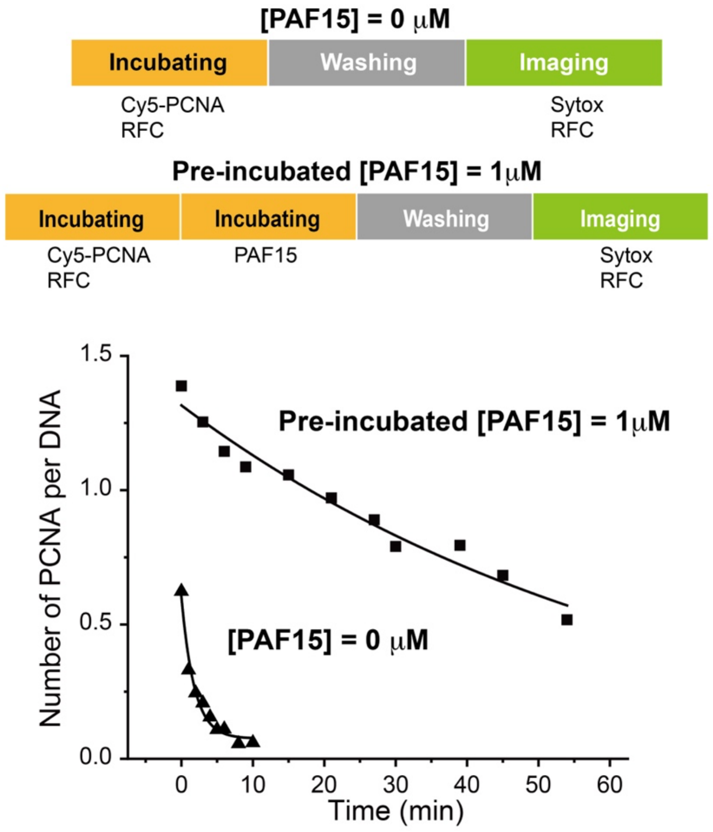


**Supplementary Figure S6. Measurement of PCNA unloading by RFC and its inhibition by bound PAF15.** PCNA dissociation from DNA was monitored over time in the presence of RFC in the imaging buffer. The number of PCNA molecules per DNA strand was quantified at each time point. The triangles represent the experiment without PAF15, while the squares represent the experiment with preincubation using 1 μM PAF15. Solid lines indicate first-order exponential decay fits for each condition. When PAF15 is not bound, PCNA dissociates significantly faster in the presence of RFC (this figure) than in its absence (half-lifetime = 1.25 ± 0.18 min; see Figure 4), consistent with RFC-mediated unloading. In contrast, when PAF15 is bound, PCNA dissociation is markedly slower (half-lifetime = 47.19 ± 30.28 min). This suggests that PAF15 stabilizes the assembly, effectively inhibiting both spontaneous dissociation and RFC-induced unloading. These results support the conclusion that the RFC-loading inhibition observed in Figure 5 is due to PAF15 bound to PCNA, rather than free PAF15 in solution.
